# Prolactin Receptor Signaling Regulates a Pregnancy-Specific Transcriptional Program in Mouse Islets

**DOI:** 10.1101/474023

**Authors:** Mark E. Pepin, Adam R. Wende, Ronadip R. Banerjee

## Abstract

Pancreatic β-cells undergo profound hyperplasia during pregnancy to maintain maternal euglycemia. Failure to reprogram β-cells into a more replicative state has been found to underlie susceptibility to gestational diabetes mellitus (GDM). We recently identified a requirement for prolactin receptor (PRLR) signaling in the metabolic adaptations to pregnancy, where mice lacking β-cell PRLR (βPRLRKO) exhibit a metabolic phenotype consistent with GDM. However, the underlying transcriptional program that is responsible for the PRLR-dependent metabolic adaptations during gestation remains incompletely understood. To identify PRLR signaling gene regulatory networks and target genes within β-cells during pregnancy, we performed a transcriptomic analysis of pancreatic islets isolated from either βPRLRKO mice or littermate controls in late gestation. Gene set enrichment analysis identified Forkhead box protein M1 (*Foxm*1) and polycomb repressor complex 2 (PRC2) subunits, *Suz*12 and *Ezh*2, as novel candidate regulators of PRLR-dependent β-cell adaptation. GO-term pathway enrichment revealed both established and novel PRLR signaling target genes that together describe a state of increased cellular metabolism and/or proliferation. In contrast to the requirement for β-cell PRLR signaling in maintaining euglycemia during pregnancy, PRLR target genes were not induced following high-fat-diet feeding. Altogether, the current study expands our understanding of which transcriptional regulators and networks mediate gene expression required for islet adaptation during pregnancy. The current work also supports the presence of pregnancy-specific adaptive mechanisms distinct from those activated by nutritional stress.

## Introduction

Gestational diabetes mellitus (GDM) is a metabolic disorder that emerges only during pregnancy, yet it presents serious health outcomes to mother and offspring even decades after birth (1). As with other forms of diabetes mellitus, a relative deficiency of functional β-cells is a major factor in the pathogenesis of GDM (2). Controversy exists, however, regarding the underlying etiology of GDM when considering its relationship with obesity and nutrition. While glycemic management of GDM can be achieved by nutritional therapy (3), only approximately half of the cases can be attributed to the metabolic effects of obesity (4,5). Furthermore, the risk of both recurrent GDM and type 2 diabetes mellitus is exceedingly high and occurs independently of obesity (6).

Unlike various other metabolic stressors that stimulate β-cell expansion, the physiologic adaptation to pregnancy comprises both an interaction between the mother and the feto-placental unit, and a rapidly fluctuating hormonal milieu that induces stereotypic changes in systemic metabolism spanning gestation and postpartum (2). While the proliferative phenotype and augmentation of β-cells seen throughout pregnancy are rapidly reversed postpartum, the adverse consequences of disturbed maternal glucose homeostasis impact both mother and developing offspring during pregnancy and long-term (1,7). These unique features of pregnancy and GDM form the premise for additional studies into the genetic relationship between GDM and type 2 diabetes and their shared or distinct pathophysiologic mechanisms (8).

Amongst the diverse extracellular and intracellular mechanisms known to contribute to gestational β-cell adaptation, the lactogens are perhaps the most well-studied (9). Most mammals, including rodents and primates, produce pituitary-derived prolactin and placental lactogens, which both signal through the prolactin receptor (PRLR) (9–12). We and others have established a critical role for PRLR signaling in rodent gestational β-cell proliferation (13), with the lack of PRLR expression in β-cells resulting in a failure of islets to proliferate during gestation (14). Conversely, β-cell specific overexpression of placental lactogens increases pancreatic islet proliferation and mass (15). Together these studies illustrate that increases in circulating lactogens during pregnancy are essential and sufficient to orchestrate key components of metabolic adaptations to pregnancy. Downstream targets of PRLR signaling that contribute to gestational proliferation include tryptophan hydroxylase 1 (*Tph1)*, the rate limiting enzyme for islet serotonin synthesis (16), transcription factors such as *Foxm1* (17) and *MafB* (14), osteoprotegerin (*Tnfrsf11b*), menin (*Men1*), as well as cyclins, cyclin-dependent kinases, cell cycle inhibitors and *Prlr* itself (14,16,18-20). However, a *global* assessment of how PRLR signaling regulates β-cell transcription during gestation is lacking.

To address this question, in the current study we examined how loss of PRLR signaling within β-cells alters pancreatic islet gene expression during pregnancy. Our findings identify both known and novel PRLR signaling targets and candidate upstream regulators of β-cell transcription. Additionally, we found that the transcriptomic program driven by PRLR signaling and its target genes is distinct from the compensatory response of islets to high-fat diet feeding, indicating pregnancy-specificity of key downstream mediators of this pathway.

## Materials and Methods

### Mouse Strains, Husbandry, Breeding and Experimentation

Female transgenic mice lacking PRLR within pancreatic β-cells (βPRLRKO) were used in the current study, as previously developed and described (14). For gestational studies, 8-week-old virgin females were mated with C57Bl6/J males and vaginal plugs scored as GD0.5. Plugged females were single-housed for the duration of pregnancy. For nutritional studies, mice were maintained on standard chow (4.7 kcal%, #7917, Envigo, Indianapolis, IN), then given *ad libitum* access to standard chow or high-fat diet (HFD, 58.0 kcal% fat, #D12492 Research Diets, New Brunswick, NJ) for either 4 or 12 weeks. Mice were group-housed (5/cage) on a 12:12-h light-dark cycle at 25 ± 1°C and constant humidity with free access to food and water except as noted. All procedures involving mice were approved and conducted in accordance with the University of Alabama at Birmingham Institutional Animal Care and Use Committee (IACUC).

### Glucose Tolerance Testing

Glucose tolerance tests (GTTs) were performed by intraperitoneal injection of glucose (1 g/kg (Chow-fed) or 2 g/kg (HF-fed) body weight, (glucose stock solution of 20% wt/vol D-glucose [Sigma] in 0.9% saline) to 16-hour overnight fasted mice. Blood glucose was determined using Bayer Contour Glucometer.

### Islet Isolation and Culture

Islets were isolated from donor mice using retrograde perfusion of the pancreatic duct with collagenase and purified using density centrifugation as previously described (14). Donor mice included βPRLRKO and littermate controls or C57Bl6/J mice either from virgin females (nonpregnant) or at gestational day (GD) 16.5 (pregnant). For in vitro experiments, hand-picked islets were cultured overnight in RPMI supplemented with 10% FBS. Subsequently, islets from individual mice were picked and divided equally into fresh media supplemented with recombinant mouse prolactin at a final concentration of 500 ng/ml (R&D Systems, Minneapolis, MN), or an equal volume of vehicle. Culture media was changed daily.

### Microarray Analysis

Microarray-based mRNA quantification was performed using the Affymetrix GeneChip Mouse 2.0 Array (ThermoFisher, Waltham, MA US). Hybridization and microarray scanning were performed by the Stanford Functional Genomics Facility (Stanford, CA). All microarray data have been deposited within the Gene Expression Omnibus repository (GSE118134). Subsequent microarray annotation, normalization, and differential expression analysis were completed in the R statistical computing environment (version 3.4.3). Briefly, raw microarray data across all samples were compiled and corrected using the robust multi-array average (RMA) of background-adjusted and quantile-normalized microarray expression data using R package *Oligo* (1.42.0) as previously described (21). Gene annotations were then generated using the MoGene-2.0 Transcript Cluster (mogene20sttranscriptcluster.db). Once expression data were examined for RNA quality, the R package *limma* (version 3.34.9) was used to generate differential gene expression using generalized linear modelling (KO versus WT) with a Benjamini-Hochberg (B-H) *P*-value adjustment (22). Due to sample size limitations (*n* = 3), dispersion estimates were first determined via maximum likelihood, assuming that genes of similar average expression strength possess similar dispersion, as previously described (23). Gene-wise dispersion estimates were then shrunken according to the empirical Bayes approach, providing normalized count data for genes proportional to both the dispersion and sample size. Differential expression was represented as log_2_ (fold-change) for each annotated gene. Statistical significance was estimated at B-H adjusted *P-*value (*Q-*value) < 0.05.

### Bioinformatics and Data Visualization

Details of the R coding scripts and other bioinformatics tools used in the current study are available on GitHub for public reference as supplemental materials (https://github.com/mepepin/bPRLRKO_Pepin.2018). Functional gene set enrichment analyses (GSEA) were performed using the interactive web-based platform *Enrichr* (24) using the Kyoto Encyclopedia of Genes and Genomes (KEGG) pathway database. Gene network analysis and visualization were performed using Cytoscape (3.7.0), with literature-curated pathway enrichment was achieved using QIAGEN’s Ingenuity Pathway Analysis (IPA^®^, QIAGEN Redwood City, www.qiagen/ingenuity) on differentially expressed genes (DEGs) using a low-stringency statistical threshold of *P* < 0.05. Heatmap and hierarchical clustering generation was performed using *pheatmap* package (1.0.8) within R, and VennPlex (24) was used to create the Venn diagrams and determine overlapping genes.

### RNA Isolation for Microarray and Quantitative Real-Time PCR Studies

Total RNA was isolated using TRIzol and reverse transcribed using the Ambion Retroscript Kit according to manufacturers’ instructions. Differential expression by quantitative PCR (qPCR) were performed with FastStart Universal Probe Master (Roche) and Thermofisher TaqMan™ assays (Waltham, MA, Supplemental Table S1 (25)) on BioRad CFX96 Touch™ (Hercules, CA) machine. Assays were performed in technical replicates and normalized to mouse actin (*Actb*) as a reference standard.

### Statistics

To determine the significance in overlap between the Venn Diagram of differentially-expressed genes identified in our study and those of previous publications, we used a hypergeometric probability distribution with Bonferroni correction to generate an adjusted *P*-value. For all targets, an unpaired two-tailed student’s t-test was performed using Tukey’s correction for multiple comparisons. Statistical significance was concluded based on a false discovery-adjusted *P*-value (*Q*) < 0.05. Functional and network GSEA, along with curated literature-supported candidate upstream regulators, were performed using QIAGEN’s Ingenuity Pathway Analysis (IPA®, QIAGEN Redwood City) unless otherwise specified. GTTs were considered significant for repeated measures ANOVA of *P*-value < 0.05.

## Results

### Differential gene expression in pancreatic islets from βPRLRKO mice during gestation

To identify genes regulated by prolactin receptor (PRLR) signaling in maternal pancreatic islets during pregnancy, we examined global transcriptomic profiles at late gestation using commercial oligonucleotide microarrays (as described in Methods). RNA purified from pancreatic islets isolated from gestational day (GD) 16.5 βPRLRKO mice (*Prlr*^f/f^; *RIP-Cre*) and littermate controls (*Prlr*^f/f^ and *Prlr*^f/+^) identified 2,542 differentially-expressed genes (DEGs) at P < 0.05; of these, 70 reached a Bonferroni-adjusted *P*-value (Q) < 0.05 (Supplemental Table S2 see (25)). Unsupervised principal components analysis (PCA) revealed that the two eigenvectors most responsible for sample variance (51.7%) were sufficient to describe a separation between βPRLRKO mice and controls, supporting that prolactin receptor knockout confers a true and measurable shift in global gene expression (Supplemental Figure S1; (26)). Gene set enrichment analysis identified Tryptophan/Serotonin metabolism and Prolactin signaling as enriched pathways (Fig. 1a), both of which are associated with the metabolic adaptation to pregnancy (13,16,27,28). We then compared the DEGs reaching Q-value significance with published studies reporting DEGs in murine islets during pregnancy in wild-type mice. We incorporated microarray data of islet gene expression during pregnancy compared to nonpregnant controls (Schraenen *et al.*; GD9.5 (29), Layden *et al.;* GD12.5. (30), and Rieck *et al.* GD14.5 (31)), and generated a Venn diagram of these studies with the 70 DEGs found in βPRLRKO mice. This comparison revealed a significant overlap of 26 DEGs (hypergeometric *P* = 2.8 × 10^−61^) (Fig. 1b). Amongst these were genes responsible for enriching pathways in Fig. 1a including tryptophan metabolism (*Tph1, Tph2*), and prolactin receptor signaling (*Prlr*), as well as diverse cellular physiologic processes (Table 1 and (25)).

**Figure 1.**
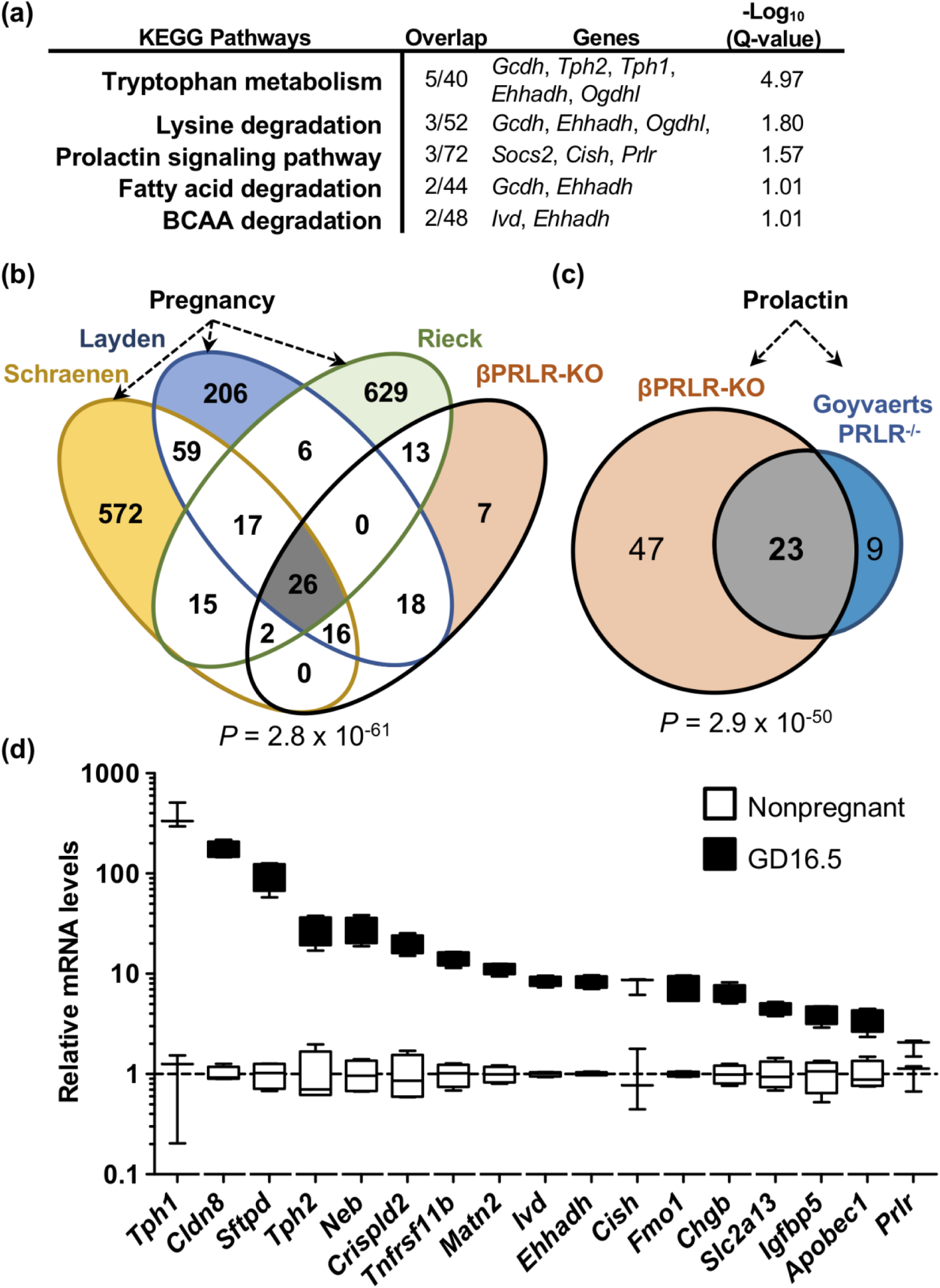
Identifying the prolactin-dependent transcriptional effects of pregnancy. **(a)** KEGG pathway enrichment using differentially-expressed genes (DEGs)* of isolated pancreatic islets from pregnant β-cell-specific prolactin receptor knockout (βPRLRKO) mice relative to littermate controls (*n* = 3). (**b)** Comparison of genes differentially regulated in pregnant βPRLRKO mice to published studies of islets from pregnant females at various stages: Schraenen *et al.* (PMID: 20886204), Layden *et al.* (PMID: 20847227), Rieck *et al.* (PMID: 19574445), and the current study (βPRLRKO). (**c)** Cross-comparison of βPRLR-dependent DEGs* identified in the current study with those identified by Goyvaerts *et al.* (PMID: 25816302) using a constitutive whole-body PRLR knockout mouse model (Q < 0.05). (**d)** qPCR validation of genes found to be induced by pregnancy and suppressed in βPRLR-KO, reported as average relative fold-change ± SEM. *Significance was assumed using Bonferroni-adjusted *P*-value < 0.05.

**Table 1.** Genes induced by pregnancy which require β-cell PRLR signaling.

| **Gene** | **Description** | **Log2**<br>**(Fold-Change)** | **q-val.** | **β-cell**<br>**physiology** |
| --- | --- | --- | --- | --- |
| *Cldn8* | claudin 8 | -4.7 | 0.0009 |  |
| *Ivd* | isovaleryl coenzyme A dehydrogenase | -2.4 | 0.002 | (32) |
| *Tph1* | tryptophan hydroxylase 1 | -5.3 | 0.002 | (14,16,33) |
| *Matn2* | matrilin 2 | -3.1 | 0.002 | (32) |
| *Cish* | cytokine inducible SH2-containing protein | -2.4 | 0.004 | (34) |
| *Tph2* | tryptophan hydroxylase 2 | -2.6 | 0.004 | (16) |
| *Socs2* | suppressor of cytokine signaling 2 | -1.7 | 0.005 | (35) |
| *Igfbp5* | insulin-like growth factor binding protein 5 | -2.6 | 0.005 | (36) |
| *B3galnt1* | UDP-GalNAc | -2.5 | 0.008 |  |
| *Slc2a13* | solute carrier family 2 (facilitated glucose transporter) | -1.4 | 0.008 |  |
| *Apobec1* | apolipoprotein B mRNA editing enzyme, catalytic polypeptide 1 | -1.6 | 0.011 |  |
| *Znrf2* | zinc and ring finger 2 | -1.3 | 0.013 |  |
| *Tnfrsf11b* | tumor necrosis factor receptor superfamily | -2.2 | 0.019 | (37) |
| *Fmo1* | flavin containing monooxygenase 1 | -1.8 | 0.021 |  |
| *Slc40a1* | solute carrier family 40 (iron-regulated transporter) | -1.5 | 0.021 |  |
| *Ehhadh* | enoyl-Coenzyme A, hydratase/3-hydroxyacyl Coenzyme A dehydrogenase | -1.8 | 0.023 | (32) |
| *Prlr* | prolactin receptor | -1.8 | 0.031 | (14,16,38) |
| *Car15* | carbonic anhydrase 15 | -1.6 | 0.033 |  |
| *Enpp2* | ectonucleotide pyrophosphatase phosphodiesterase 2 | -1.1 | 0.033 |  |
| *Igfals* | insulin-like growth factor binding protein, acid labile subunit | -0.9 | 0.033 |  |
| *Gcdh* | glutaryl-Coenzyme A dehydrogenase | -0.9 | 0.041 |  |
| *Wipi1* | WD repeat domain, phosphoinositide interacting 1 | -1.3 | 0.043 |  |
| *Aqp4* | aquaporin 4 | -2.1 | 0.045 |  |
| *Chgb* | chromogranin B | -1.4 | 0.046 | (32) |
| *Slc6a8* | solute carrier family 6 (neurotransmitter transporter, creatine) | -1.0 | 0.047 |  |
| *Gnb1l* | guanine nucleotide binding protein (G protein), beta polypeptide 1-like | -0.9 | 0.049 |  |

Thus, this approach identified 26 genes that are induced by pregnancy in a PRLR-dependent manner. We also mined microarray data of islet gene expression in global PRLR-knockout females (*Prlr^−/−^*; created by (39)) examined by Goyvaerts *et al.* at early gestation (GD9.5) (40). Although the global *Prlr^−/−^* and βPRLRKO models differ in several key respects (9,14,39,41), this cross-model comparison showed a significant overlap (hypergeometric *P* = 2.9 × 10^−50^) in differential gene expression (defined as Q < 0.05) (Fig. 1c). To further validate our microarray findings, we examined gene expression using qPCR on an independent set of wild-type GD16.5 and nonpregnant C57Bl6/J female mice, which revealed significant changes in expression consistent with array data (Fig. 1d). Altogether, these results indicate that the effects of PRLR signaling on β-cell gene activation occur independently of PRLR signaling in other tissues, and are present at both early and late gestation.

### Prolactin and high-fat feeding stimulate divergent transcriptional cascades in the β-cell

Our analysis establishes that PRLR signaling is necessary for the induction of a subset of genes in islets during healthy gestation. However, PRLR signaling may itself interact with the hormonal milieu of pregnancy to regulate transcription. To determine whether PRLR signaling is sufficient to regulate genes in an islet-autonomous fashion, we treated cultured islets for 24 hours with recombinant prolactin. This treatment significantly upregulated many of the PRLR-dependent DEGs (Fig. 2a), including *Tph1* and *Tph2* as previously reported (29). Our findings therefore demonstrate that PRLR signaling is both sufficient and required to regulate a subset of DEGs normally induced by pregnancy.

**Figure 2.**
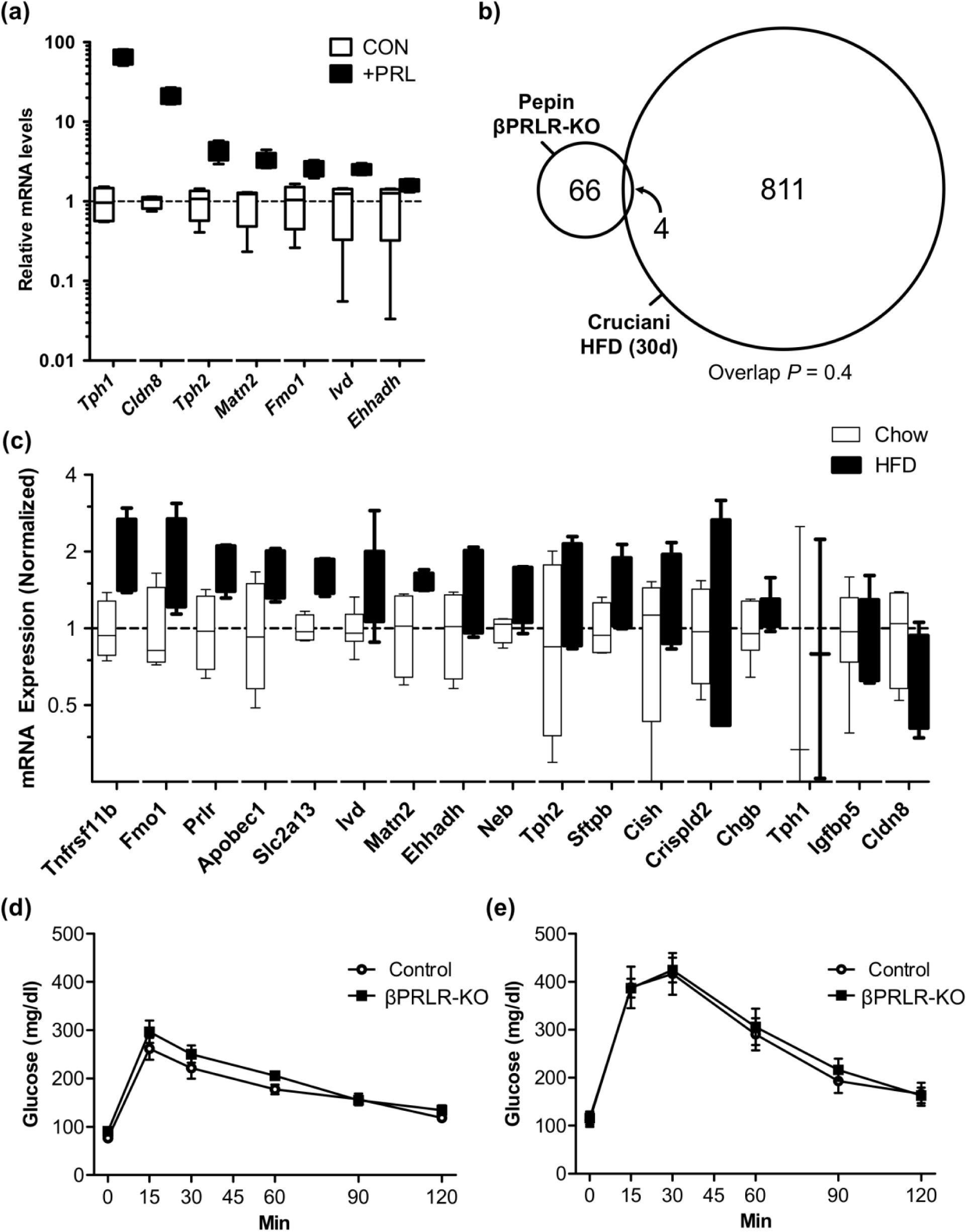
Pregnancy and high-fat diet feeding activate divergent transcriptional programs. **(a)** qPCR quantification of PRLR-dependent pregnancy genes in primary isolated pancreatic islets treated with/without 500 ng/ml recombinant mouse prolactin (PRL+/−) for 24 hours (*n* = 4). (**b)** Venn diagram illustrating the degree of overlap between βPRLR-dependent DEGs and those induced by high-fat diet (HFD) based on RNA-sequencing dataset generated by Cruciani et al. (PMID: 28377873). **(c)** qPCR quantification of PRLR-dependent pregnancy genes in primary isolated pancreatic islets from chow-or HFD-fed mice (*n* = 4). **(d)** Intraperitoneal glucose tolerance testing of βPRLRKO mice and littermate controls at baseline **(e)** following four weeks of standard chow (Chow) or 60% HFD feeding (*n* = 6). *Significance of transcriptomic data was assumed using Bonferroni-adjusted *P*-value < 0.05.

Metabolic stressors including nutrition and pregnancy have been theorized to converge on the β-cell, leading to its dysfunction and ultimately diabetes mellitus; however, empirical evidence supporting this mechanistic convergence remain inconclusive (1,42). Because pregnancy initiates an adaptive metabolic response via profound increases in lactogenic and other hormones, we hypothesized that PRLR-mediated adaptations activate a pregnancy-specific adaptive mechanism in pancreatic islets. To test this, we first compared our βPRLRKO dataset to published RNA-sequencing data of gene expression changes following high-fat diet (HFD), a widely-used metabolic stress model system (43) (Fig. 2b). Surprisingly, the overlap of DEGs was minimal and not significant (Hypergeometric *P* = 0.4), comprising only 4 genes (*Matn2*, *2410021H03Rik*, *Igfbp5*, *Cntn3*) that were differentially co-expressed by both βPRLRKO and 30-day HFD feeding. As Cruciani *et al.* only examined male mice (43), we performed a 4-week nutritional study using female mice to determine whether high-fat diet (HFD) feeding displayed sexually dimorphic effects on the 70 DEGs reaching Q-value significance in βPRLRKOs. Consistent with findings in male mice, females exhibited no significant changes in these genes following 4 weeks of high-fat diet (Fig. 2c). Thus, for both sexes, HFD feeding was incapable of inducing PRLR-dependent DEGs.

We next sought to determine whether the PRLR is required for β-cell physiologic adaptation to HFD feeding. Unlike the requirement for intact PRLR signaling in maintaining gestational glucose homeostasis, both βPRLRKO female mice and littermate controls had no baseline difference in GTT (Fig. 2d) and developed equivalent glucose intolerance following four weeks of HFD feeding (Fig. 2e). These results support our hypothesis that both transcriptional regulation and physiologic adaptation of β-cells by PRLR signaling employs pregnancy-specific mechanisms that differ from those of HFD adaptation.

### Pathway analysis of differential gene expression

To identify potential mechanisms underlying pregnancy-specific adaptations in islets, we next sought to understand the global transcriptional landscape that is regulated by PRLR signaling in pregnancy. Because the FDR-adjusted threshold yielded only 70 DEGs, we considered whether reducing the statistical stringency of our analysis (*P* < 0.05) would capture nodal networks and regulatory clusters. Hierarchical clustering and heatmap visualization justified this approach, displaying a distinct expression profile even at this cutoff (Fig. 3a). KEGG Pathway enrichment of the 2,540 genes (*P* < 0.05) revealed a clear signature of cellular proliferation amongst all of the top-5 most enriched networks: G2/M DNA Damage Checkpoint (*P* = 1.7 × 10^−5^), GADD45 Signaling (*P* = 5.6 × 10^−5^), Regulation of Cellular Mechanics (*P* = 2.3 × 10^−4^), Estrogen-mediated S-phase Entry (*P* = 2.8 × 10^−4^), and Mitotic Roles of Polo-Like Kinase (*P* = 4.9 × 10^−4^). Volcano plot revealed a negatively skewed signature of transcriptional changes by βPRLRKO relative to littermate controls (Fig. 3b), consistent with known upregulation of β-cell genes during pregnancy (30,31).

**Figure 3.**
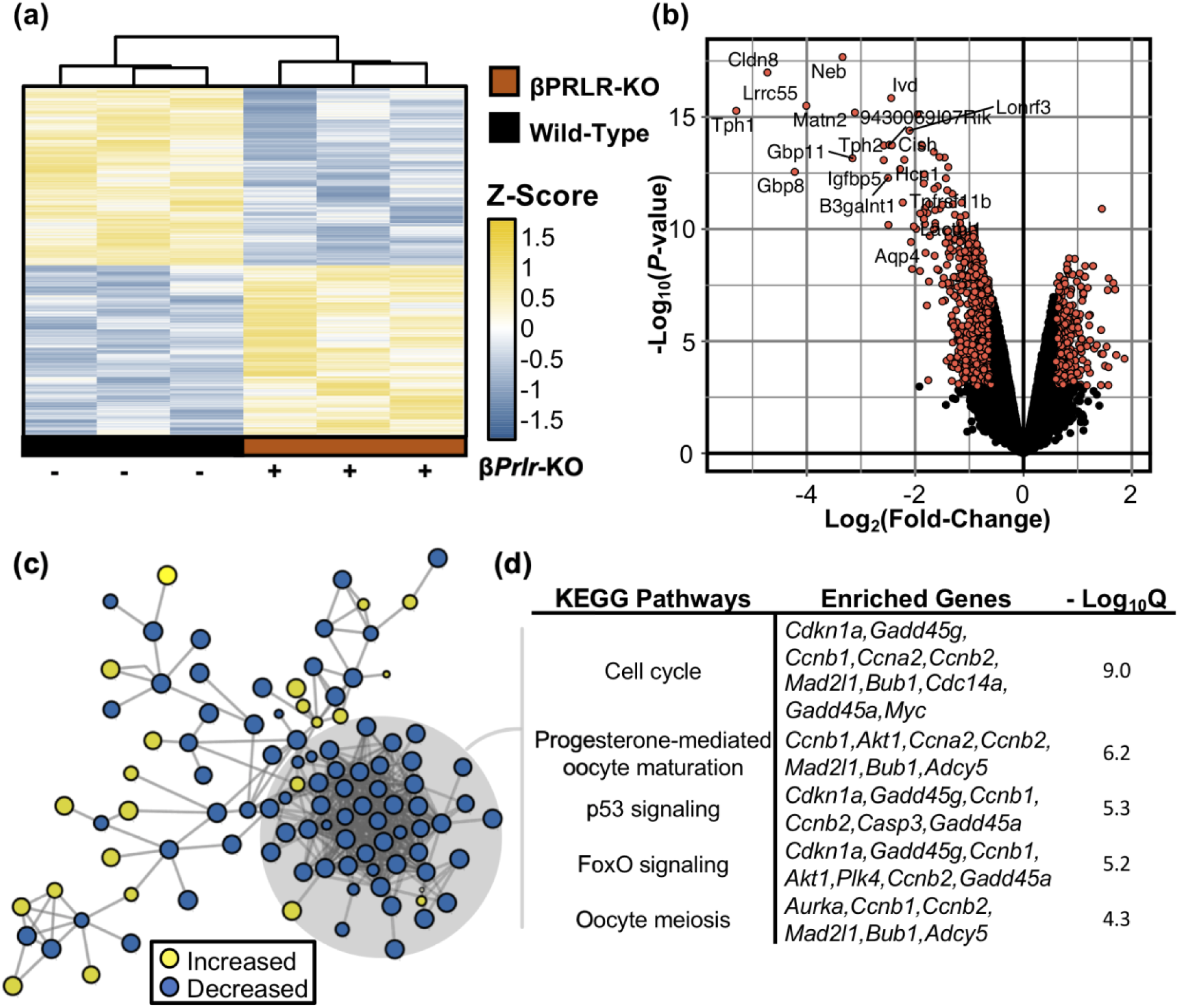
Gene expression analysis of βPRLRKO identifies proliferative impairment. **(a)** Hierarchical clustering of normalized beta values via Wald.D2 test and dendrogram constructed by Euclidean distance with heatmap visualization of DEGs*. **(b)** Volcano plot illustrating the distribution of genes based on -Log_10_(P-value) as a function of Log_10_(Fold—Change), labelling the most differentially-regulated genes (|Fold-change| > 2, FDR < 0.05) **(c)** Protein-protein interaction network generated from DEGs* based on the STRING database. **(d)** Gene set enrichment of the gene module with highest degree of protein interactivity using KEGG pathways of DEGs. *significance threshold of *P* < 0.05 was used for gene expression, followed by BH-adjusted *P* (“Q”) < 0.05 for subsequent gene set enrichment analysis.

Our gene set enrichment analysis identified established pathways differentially affected by loss of PRLR signaling. However, to identify novel networks affected in βPRLRKO, we used the STRING database (44) to generate a network of DEGs based on known and predicted interactions among their encoded proteins (Fig. 3c). After ranking nodes by degree of interaction, the largest cluster was identified to functionally enrich pathways associated with cellular proliferation and developmental programs (Fig. 3d): cell cycle (BH-adjusted *P* = 10^−9^), progesterone-mediated oocyte maturation (BH-adjusted *P* = 10^−6.2^), p53 signaling (BH-adjusted *P* = 10^−5.3^), FoxO signaling (10^−5.2^), and oocyte meiosis (BH-adjusted *P* = 10^−4.3^). By contrast, when protein-protein interaction analysis was performed on the dataset from mice fed high-fat diet (43), the network instead enriched pathways mediating fatty acid metabolism and signaling, with peroxisome proliferator-activated receptors (PPARs) as the most-enriched transcriptional regulators (Supplemental Figure S2; (26)). These data demonstrate that βPRLRKO impairs a pregnancy-specific program of β-cell proliferation during pregnancy.

### EZH2/PRC2 and FOXM1 are PRLR-dependent candidate transcriptional regulators of β-cell adaptation in pregnancy

To identify upstream regulators capable of driving this proliferative signature, we used the Ingenuity^®^ (IPA) causal network analysis to leverage the directionality of gene changes when performing gene set enrichment (45). IPA upstream functional analysis disproportionately enriched genes known to interact with PRLR itself, both directly and indirectly (Fig. 4a). Other upstream regulators included Growth and DNA Damage Inducible 45 γ (*Gadd45g*, 2.1-fold suppressed) and DNA methyltransferase 3B (*Dnmt3b*, 2.2-fold suppressed), both regulators of *de novo* DNA methylation (46,47). Examination of the genes reported to interact with PRLR revealed increased expression of upstream factors and decreased expression of PRLR target genes (Fig. 4b). Of the PRLR target genes, we noted that the histone methyltransferase Enhancer of Zeste Homologue 2 (*Ezh2*) was suppressed in βPRLRKO.

**Figure 4.**
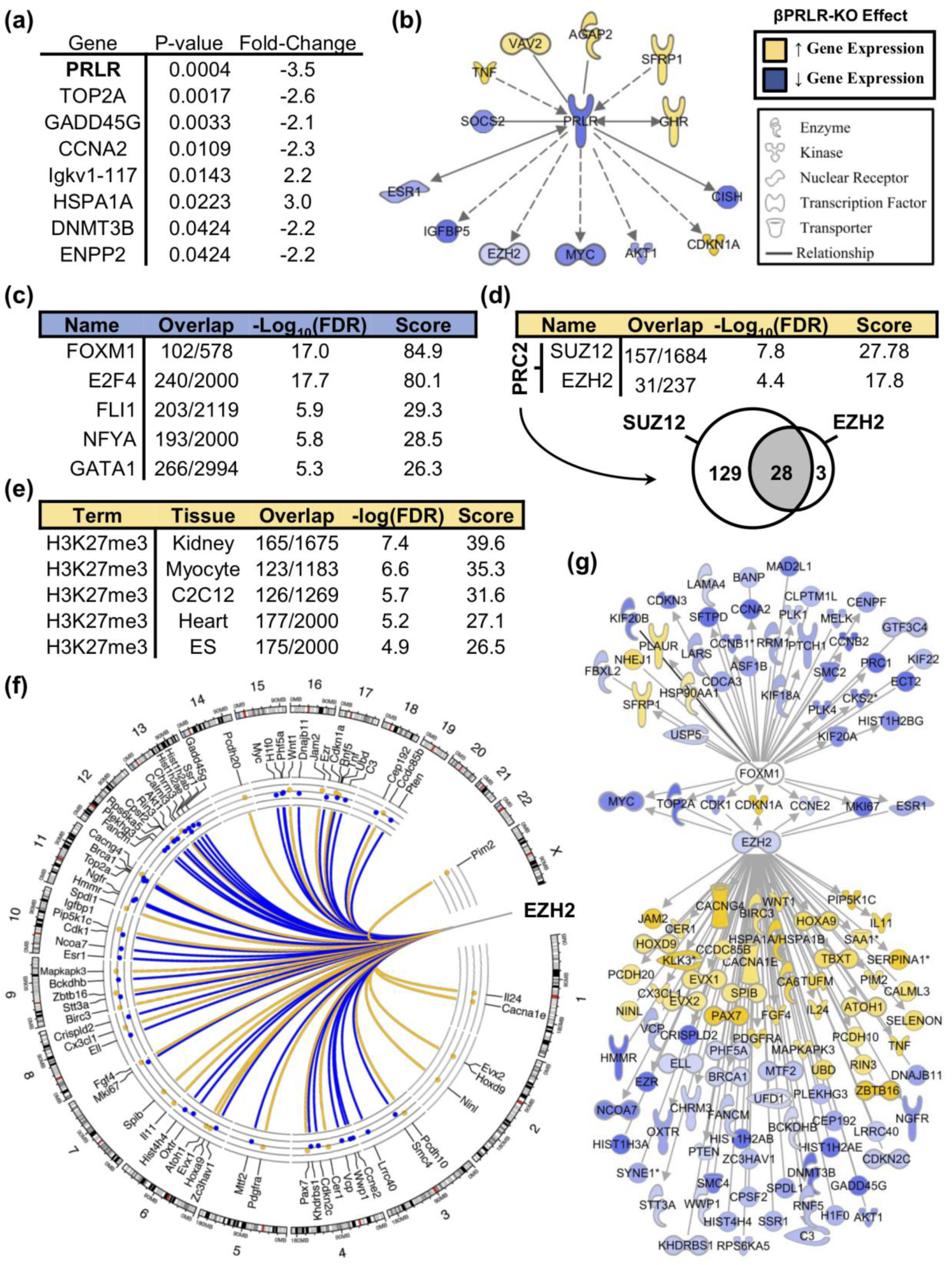
EZH2 and Foxm1 as likely upstream regulators of PRLR-dependent gene expression. **(a)** PRLR identified as top differentially-expressed upstream regulator via Ingenuity database (IPA^®^ Redwood City, CA). **(b)** Regulatory network of DEGs* associated with PRLR (**blue** = down-regulated; **yellow** = up-regulated). \**P* < 0.05 for pathway and network enrichment. **(c)** Upstream DNA-binding protein enrichment of up-regulated differentially-expressed genes (DEGs) and **(d)** down-regulated DEGs in βPRLRKO relative to WT was performed using published ChIP-Sequencing datasets based on the Encyclopedia of DNA Elements (ENCODE). **(e)** Up-regulated DEGs were enriched for histone modifications using ENCODE database. **(f)** Circular genome plot of differentially-regulated direct downstream targets of EZH2. **(g)** Co-regulatory network of downstream DEGs of EZH2-FOXM1 regulation.

As a transmembrane receptor resident at the cell surface, PRLR indirectly regulates transcription through signal transduction cascades, including the canonical JAK2-STAT5 pathway (48). However, as mice with a β-cell specific deletion of Stat5 do not exhibit defective glucose homeostasis or GDM during pregnancy (49), we hypothesized that multiple transcription factors coordinately regulate transcription for PRLR signaling-mediated gestational adaptations. To identify candidates, we inspected the proximal promoter region (−1kB to +500kB) of differentially-regulated genes (*P* < 0.05) for disproportionate enrichment of transcription factor binding sites.

Using the ENCODE ChIP-sequencing database (50), we identified promoter binding sites for *Foxm1* and *E2f4* among down-regulated DEGs (Fig. 4c), a finding consistent with prior studies (14,17). *Stat5a* was also statistically enriched (FDR < 0.05), though not among the 10 most-enriched putative regulators. Conversely, promoter enrichment of genes expressed at higher levels in βPRLRKO identified promoter binding sites for *Suz12* (BH-adjusted *P* = 10^−7.8^) and *Ezh2* (BH-adjusted *P* = 10^−4.4^), two components of the polycomb repressor complex (PRC2) (Fig. 4d). Co-localization of *Ezh2* and *Suz12* promoter binding sites DEGs was also noted, demonstrating a 90% (28/31) overlap. To determine whether PRC2 methyltransferase activity likely controls gene expression in βPRLRKO, we then enriched DEG promoters using ChIP-sequencing analysis of chromatin modifications using the human histone modification database (51). Histone modification enrichment identified H3K27 trimethylation (H3K27me3), the modification enzymatically mediated by EZH2/PRC2, as the only histone modification localized to up-regulated DEG promoters (Fig. 4e). Reciprocally, the promoter regions of down-regulated DEGs contained sites of H3K27 acetylation (H3K27ac), the counter-regulatory modification of H3K27me3. Consistent with other research (52,53), these observations support that the PRC2-mediated gene silencing that occurs during pregnancy is de-repressed in the setting of βPRLRKO.

*Ezh2* has been classically identified as a transcriptional repressor via histone-methyltransferase activity in the PRC2 (54). However, we and others have identified a paradoxical role for *Ezh2* as a positive regulator of gene expression upon heterodimerization with *Foxm1* and other transcriptional regulators (55–57). Therefore, to understand whether this dual role of *Ezh2* might be simultaneously present in pancreatic islets, we plotted the 75 DEGs (*P* < 0.05) that are targets of *Ezh2* according to the Ingenuity^®^ database (Fig. 4f). The resulting distribution depicted a broad array of target genes both increased and decreased by βPRLRKO. Additionally, genes co-targeted by *Ezh2* and *Foxm1* feature key regulators of cellular proliferation (Fig. 4g): *Top2a* (log_2_fold = −1.4, *P* = 0.001), *Mki67* (log_2_fold = −1.4, *P* = 0.001), *Ccne2* (log_2_fold = −0.30, *P* = 0.04), *Myc* (log_2_fold = −0.81, *P* = 0.0007), *Esr1* (log_2_fold = −0.60, *P* = 0.005), *Cdk1* (log_2_fold = −0.6, *P* = 0.001), and *Cdkn1a* (log_2_fold = 0.87, *P* = 0.0096). In sum, our analyses support a role of *Foxm1* and *Ezh2* as co-regulators of the proliferative signature within β-cells during pregnancy.

## Discussion

Prior studies have identified transcriptional changes within pancreatic islets during pregnancy, a period of rapid β-cell proliferation and mass expansion (29–31). Many hormonal and metabolic changes accompany gestation, with lactogens serving a critical role in physiologic adaptation of β-cells (14,15). While the mitogenic effects of lactogen signaling have been widely studied, the specific gene targets and transcriptional networks under lactogen control remain poorly understood. Using a β-cell specific prolactin receptor knockout (βPRLRKO) mouse model, we confirm the requirement of intact β-cell PRLR signaling for induction of both well-established and novel genes and gene networks during pregnancy.

We identified and cross-validated a number of known and novel pro-proliferative genes as induced by pregnancy in a PRLR-dependent manner (Table 1). Our findings are consistent with a recent proteomic analysis of islets during pregnancy, which independently found proteins encoded by our DEGs to be differentially regulated in late gestation: *Ivd, Matn2, Ehhadh, Myc*, and *Chgb* (32). Amongst these, we confirmed the requirement of intact β-cell PRLR signaling for induction of well-established gestational pro-proliferative genes (e.g. *Tph1*, *Tph2*, *Prlr*, *Opg*). In addition to previously identified genes of pregnancy, we identified many novel gene targets of PRLR signaling that provide exciting avenues for future study into the mechanisms of islet gestational adaptation. For instance, pregnancy increases gap junction coupling between β-cells (11), which has been theorized to regulate insulin secretion and the alterations in insulin secretion during pregnancy (58). Claudin-8 (*Cldn8*), here identified as induced by PRLR in pregnancy, is an integral membrane protein and tight junction component that has not been studied in β-cells in any context.

In our search for regulatory networks impacted by PRLR disruption in pregnancy, we identified *Foxm1* and *Ezh2* as putative nodal upstream co-regulators of gestational gene expression and cellular proliferation (Fig. 5). Within the pancreatic islet, *Ezh2* has previously been shown to silence the tumor suppressor Ink4a/Arf (p16) in aging and diabetes to increase β-cell proliferation (59,60). Conversely, selective β-cell *Ezh2* knockout in mice has been shown to produce hyperglycemia in mice (59). In the current study, we found that pregnancy induces *Ezh2* in a PRLR-dependent manner. Therefore, gestational induction of *Ezh2* likely creates a pro-proliferative state required for metabolic adaptation. Our results thus broaden our understanding of *Ezh2* as a regulator of β-cell function.

**Figure 5.**
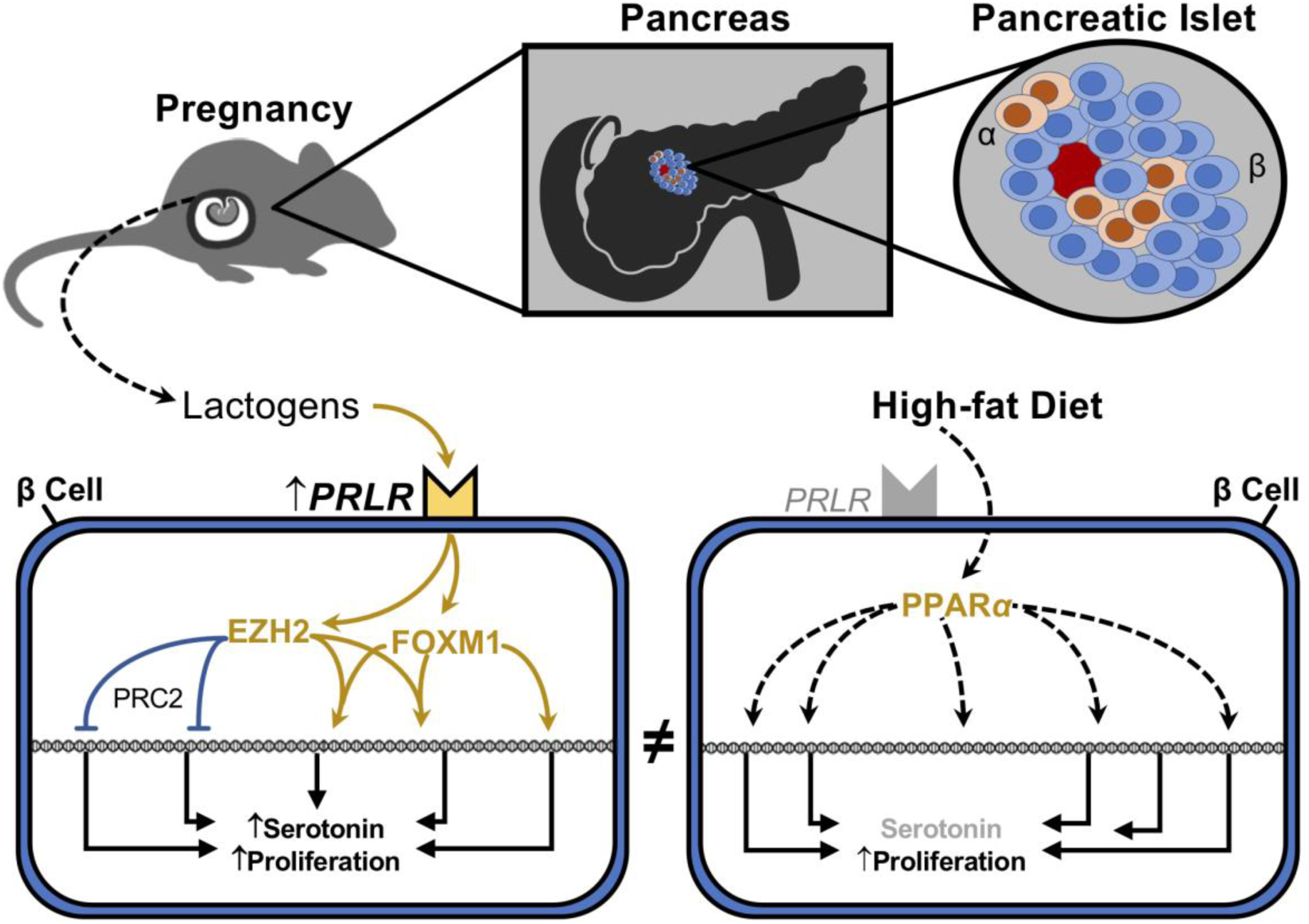
Proposed model of pregnancy-specific transcriptional reprogramming. Placental and pituitary lactogens induce transcriptional activation of serotonin signaling and proliferation within β-cells via *Ezh2* and *Foxm1*. In contrast, high-fat diet feeding induces transcriptional reprogramming via PPARα independent of the prolactin receptor and serotonin signaling.

Though canonically associated with gene silencing via histone 3 trimethylation (H3K27me3), *Ezh2* has also been shown to heterodimerize with transcription factors, such as *Foxm1*, to activate gene expression in a methylation-independent mechanism (57,61). In breast and prostate tissue, *Ezh2* is induced by PRLR activation via STAT5, and mediates the mitogenic effects of prolactin in these tissues (62,63). Our results therefore support a broader role for *Ezh2* in the regulation of β-cell proliferation during pregnancy, analogous to its role in other tissues (57).

As GDM and type 2 diabetes share many similarities and risk factors, it has been widely assumed that they share underlying pathophysiologic mechanisms. However, our data demonstrate that the reprogramming of pancreatic islets during pregnancy is distinct from that of nutritional stressors. We found that the robust PRLR-dependent transcriptional changes in gestation were not regulated by HFD feeding. Co-expression network analysis further identified distinct functional enrichment in pregnancy associated with cellular proliferation, whereas HFD feeding enriched fatty acid signaling via PPARs. Our analysis therefore provides several avenues for investigation whereby disease-specificity may be realized. Stable modifications to gene accessibility and expression, termed epigenetic changes, that we have identified as etiology-specific drivers of other diseases (57). Here, we identify many PRLR-dependent epigenetic modifiers that regulate chromatin remodeling (via *Ezh2* and *Suz12*) and/or differential DNA methylation (via *Gadd45*g and *Dnmt3b*). Therefore, the pregnancy-specific response to PRLR signaling likely requires many genetic and epigenetic regulators to coordinate β-cell adaptation. Nevertheless, future studies are needed to mechanistically define these programs.

While our findings underscore the importance of PRLR signaling as a mediator of β-cell function in pregnancy, we recognize several limitations to our studies. We and others have established that PRLR is expressed specifically in β-cells of rodent islets (14,64). However, we cannot exclude the possibility of non-β-cell effects of βPRLRKO. Therefore, future studies should employ single-cell or cell-sorted approaches to analyze gestational islet transcription. The computational approach we used to define transcriptional regulators of PRLR signaling provide preliminary support for their role in gestational adaptation; however, future mechanistic studies are needed to determine the nature of their influence. Additionally, although gestational induction of DEGs in our data is likely to be directly downstream of PRLR signaling, we recognize that parallel compensatory signaling mechanisms exist which may contribute to the differential regulation of gene expression. Consistent with this possibility, we observed a subtle increase in GHR expression on microarray analysis (1.3-fold, *P* = 0.03), a known inducer of the prolactin receptor (65).

In conclusion, the current study examines novel pregnancy-specific gene networks and PRLR-dependent transcriptional programs that describe a state of augmented cellular proliferation. Despite the stated limitations, we believe that the computational approach, combined with both *in vivo* and *in vitro* experimentation, serves to significantly advance our understanding of how β-cells respond to the metabolic demands of pregnancy. These lactogen-mediated regulatory networks likely reflect broader adaptive mechanisms that apply to other contexts. We propose PRLR-dependent transcriptional targets and upstream regulators as exciting new candidates to examine in gestational diabetes pathogenesis.

## Acknowledgements

Financial support for this work provided to R.R.B. by University of Alabama School of Medicine Start-up funds and the UAB Diabetes Research Center Pilot and Feasibility Grant (P30 DK079626) and NIH R01 HL133011 to A.R.W. Training support was provided to M.E.P. by the NIH F30 HL137240. R.R.B. designed the study. M.E.P and R.R.B performed experiments, collected data, and analyzed the results. All authors contributed to the writing and editing of the manuscript. The authors would like to thank Drs. Kirk Habegger, Stuart Frank, Chad Hunter, and Sushant Bhatnagar for helpful discussions and critical reading of the manuscript.

